# Activation of FAM111A Protease Induces Defects in Nuclear Function that Likely Underlie its Roles in Disease and Viral Restriction

**DOI:** 10.1101/2020.05.04.077594

**Authors:** Minghua Nie, Martina Oravcová, Yasaman Jami-Alahmadi, James A. Wohlschlegel, Eros Lazzerini-Denchi, Michael N. Boddy

## Abstract

Mutations in the nuclear trypsin-like serine protease FAM111A cause Kenny-Caffey syndrome (KCS2) with hypoparathyroidism and skeletal dysplasia, or perinatally lethal osteocraniostenosis (OCS). In addition, FAM111A was identified as a restriction factor for certain host range mutants of the SV40 polyomavirus and VACV orthopoxvirus. However, because FAM111A function is poorly characterized, its roles in restricting viral replication and the etiology of KCS2 and OCS remain undefined. We find that the FAM111A KCS2 and OCS patient mutants are hyperactive, inducing apoptosis-like phenotypes in a protease-dependent manner. Similarly, in response to the attempted replication of SV40 host range mutants in restrictive cells, FAM111A activity induces the loss of nuclear barrier function and structure. Interestingly, pan-caspase inhibitors do not block FAM111A-dependent phenotypes such as nuclear “leakiness”, shrinkage and pore redistribution, implying it acts independently or upstream of caspases. In this regard, we identified nucleoporins and the associated GANP transcription factor as FAM111A interactors and candidate targets. Together our data provide key insight into how FAM111A activation can restrict viral replication, and how its deregulated activity could cause KCS2 and OCS.

## Introduction

Family With Sequence Similarity 111 Member A (FAM111A) was identified as an interactor of the SV40 large T antigen (LT), and a host range restriction factor that antagonizes productive infection by certain SV40 mutants (Fine et al., 2012). Specifically, FAM111A depletion rescues replication center formation and gene expression defects of LT mutants lacking C-terminal sequences (Tarnita et al., 2019). Further analysis indicated that FAM111A executes its restrictive function during or following SV40 viral genome replication (Tarnita et al., 2019).

Expanding the role of FAM111A in protecting against viral challenge, it was identified along with the PCNA clamp-loading complex RFC1-5 as a restriction factor for orthopoxvirus host range mutants lacking the serine protease inhibitor SPI-1 (Panda et al., 2017). Early studies on rabbitpox virus (RPXV) host range mutants lacking SPI-1 suggested that restrictive human A459 cells exhibit apoptosis-like features upon infection, which required RPXV DNA synthesis (Brooks et al., 1995). However, a subsequent study found that the related vaccinia virus (VACV) lacking SPI-1 caused nuclear morphological changes such as chromatin condensation and membrane invagination, without the normal biochemical features of apoptosis (Shisler et al., 1999).

Thus, viruses with diverse life cycles e.g. nuclear replication sites (polyomavirus) versus replication in the cytoplasm (orthopoxvirus) possess a mechanism to overcome the antagonistic effects of FAM111A. However, how FAM111A combats viral challenges remains poorly characterized, but its activity is likely triggered during viral replication (Panda et al., 2017, Luttge and Moyer, 2005, Tarnita et al., 2019).

Consistent with its apparent replication-dependent role in viral restriction, FAM111A was identified in high-throughput screens as a protein enriched on nascent chromatin i.e. replication forks (Alabert et al., 2014, Wessel et al., 2019). Exogenous FAM111A was found to associate with replication sites via a PCNA interacting motif (Alabert et al., 2014), whereas endogenous FAM111A was detected in nucleoli until S phase when despite elevated expression it became undetectable (Tarnita et al., 2019).

The FAM111A C-terminus contains a predicted trypsin-like serine protease domain with an as yet untested catalytic triad. However, the role of vaccinia virus SPI-1 in antagonizing FAM111A is consistent with the presence of protease activity (Panda et al., 2017). Notably, dominant mutations in the FAM111A protease domain cause Kenny-Caffey syndrome type 2 (KCS2) and osteocraniostenosis (OCS), which present with hypoparathyroidism, short stature, and skeletal defects (Unger et al., 2013, Isojima et al., 2014, Nikkel et al., 2014). How the KCS2 and OCS mutations impact FAM111A activity remains unknown but due to their position in the domain and their dominance over wild-type they are suggested to be gain-of-function (Unger et al., 2013).

We recently identified FAM111A as a novel interactor of the SUMO-targeted ubiquitin ligase STUbL, which directs SUMO conjugated proteins for ubiquitination and extraction from chromatin or protein complexes, coupled or not to protein degradation by the proteasome (Nie and Boddy, 2016, Geoffroy and Hay, 2009, Nie et al., 2012, Nie and Boddy, 2015, Nie et al., 2017, Westerbeck et al., 2014). Whilst the STUbL-FAM111A regulatory axis will be the subject of future studies, we present data that together with recent high-throughput analyses show that FAM111A is conjugated to both SUMO1 and SUMO2 and is a likely STUbL target. We find that endogenous FAM111A colocalizes with PCNA during DNA replication, which places it amongst protein complexes regulated by SUMO modification and STUbL (Keiten-Schmitz et al., 2019).

Importantly, we show that hyperactive FAM111A is cytotoxic, inducing early apoptosis-like phenotypes that are insensitive to pan-caspase inhibitors. In particular, we show that the KCS2 and OCS FAM111A patient mutants are hyperactive and cytotoxic due to deregulated protease activity. Expression of these mutants perturbs nuclear morphology, nuclear pore distribution, DNA replication, and cell-substrate attachment, phenotypes that are insensitive to pan-caspase inhibitors.

We also find that FAM111A’s cytotoxic effects are not restricted to the patient mutants of FAM111A. That is, in response to the replication of polyomavirus (SV40) host range mutants in restrictive cells, endogenous FAM111A also disrupts nuclear barrier function, nuclear morphology and nuclear pore distribution. These effects likely explain or contribute to the restrictive activity of FAM111A during viral challenge. Interestingly in light of these FAM111A-mediated phenotypes, we identified several nuclear pore components and an associated TREX-2 mRNA export factor as candidate substrates of FAM111A.

Overall, our data provide key new insights into the pathological effects of KCS2/OCS FAM111A mutants, and the functions of endogenous FAM111A in viral restriction. In addition, the apparent caspase-independent effects of FAM111A protease activity on nuclear form and function lend credence to the controversial role of serine protease(s) upstream of caspases in triggering early alterations in the nuclear permeability barrier (Egger et al., 2003, O’Connell and Stenson-Cox, 2007, Ferrando-May et al., 2001, Strasser et al., 2012, Kopeina et al., 2018).

## Results

### FAM111A Identified by Proximity Labeling of RNF4 Proteomic Environment

We used proximity labeling (BioID) to identify potential interactors and/or substrates of the human SUMO-targeted ubiquitin ligase (STUbL) RNF4 (Dantuma and van Attikum, 2016, Nie and Boddy, 2016, Roux et al., 2012), the results of which will be published elsewhere. Here we focus on the intriguing candidate RNF4 interactor FAM111A. Of note, FAM111A was also identified in distinct high-throughput screens for RNF4 targets and SUMO2 conjugates but not characterized further (Kumar et al., 2017, Liebelt et al., 2020).

Reproducing our screen conditions confirmed that RNF4 is in close proximity to FAM111A in cells (Figure 1A). Notably, RNF4^CS12^, the ubiquitin ligase dead mutant of RNF4, in which cysteines 132, 135, 173, and 176 were replaced with serines (Tatham et al., 2008) resulted in much stronger labeling of FAM111A, as compared to wild-type. This is consistent with the role of RNF4 in promoting the turnover of SUMO-conjugated targets, including FAM111A in a high-throughput study (Dantuma and van Attikum, 2016, Geoffroy and Hay, 2009, Kumar et al., 2017).

**Figure 1.**
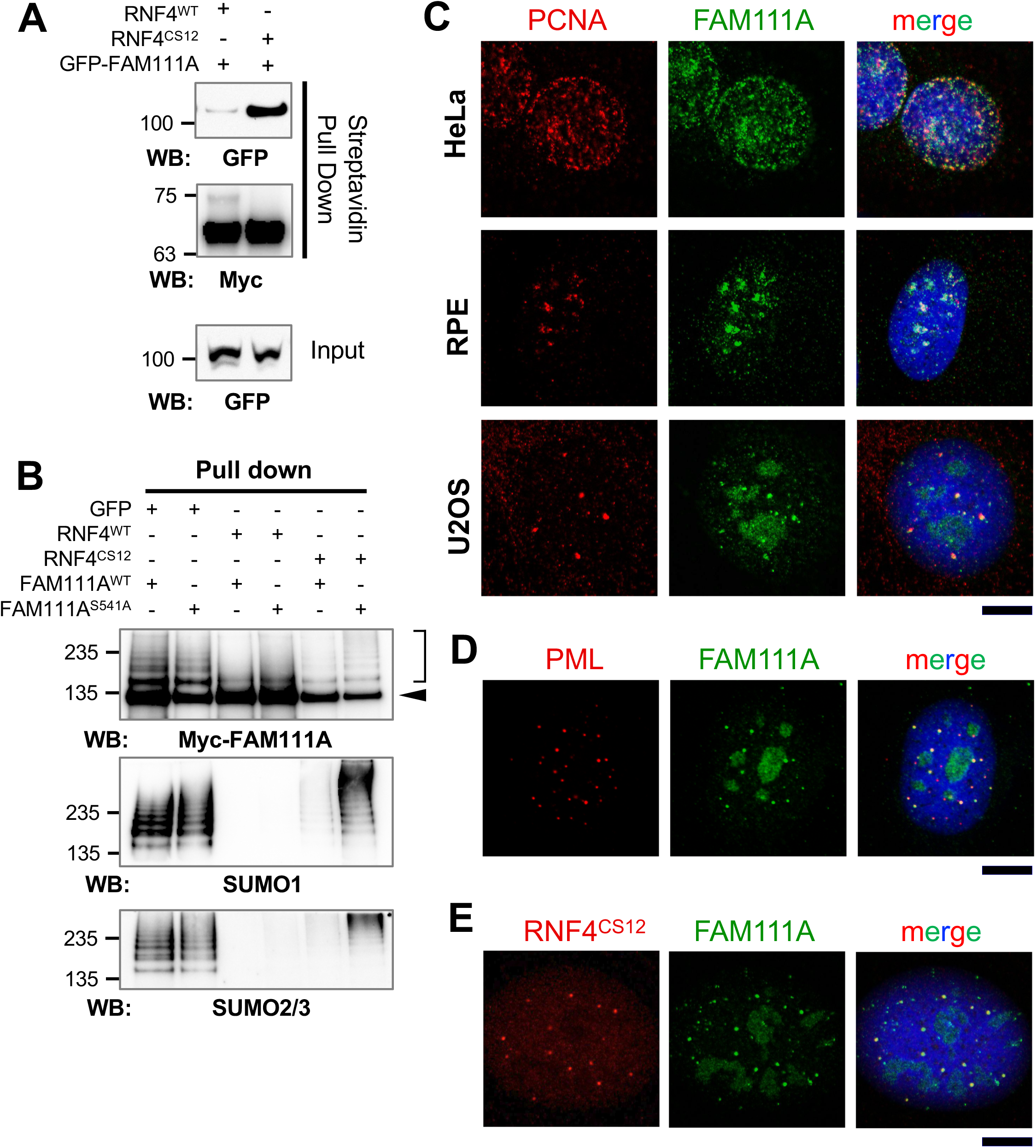
FAM111A interacts with RNF4, and is SUMOylated. Its localization and levels were characterized in multiple cell lines. **A.** HEK293T cells co-transfected with plasmids expressing MycBioID-RNF4 and GFP-FAM111A were supplied with biotin 4 h after transfection. The cells were harvested 28 h post-transfection. Equal numbers of cells from each sample were used for pulldown. Proteins bound to streptavidin magnetic beads were resolved by SDS-PAGE, and analysis by Western blotting using GFP and Myc antibodies. **B.** HEK293T cells co-transfected with plasmids expressing MycBioID-FAM111A and GFP-RNF4 or GFP along were supplied with biotin 4 h after transfection and harvested 24 h later. Pulldown with streptavidin magnetic beads and Western blotting were performed as described above, using Myc, SUMO1 and SUMO2/3 antibodies. Arrow, unmodified MycBioID-FAM111A (110 KDa); bracket, SUMO-modified FAM111A. **C.** U2OS, HeLa, RPE cell lines were pre-extracted with Triton X-100 before fixed with formaldehyde and co-stained with DAPI and antibodies for PCNA and FAM111A. **D.** U2OS cells were stained with antibodies against PML and FAM111A. **E.** U2OS cell line expressing Myc-RNF4^CS12^ under doxycycline-inducible promoter was induced to express RNF4. The cells were stained with Myc and FAM111A antibodies as well as DAPI, and then imaged by confocal microscopy. Scale bars, 10 µm (**C** to **E**).

We next tested the SUMOylation state of FAM111A and how it is impacted by RNF4. We expressed FAM111A together with wild-type RNF4, RNF4^CS12^, or a GFP control. As can be seen, purification and western analysis of FAM111A under denaturing conditions reveals a band of the expected size for FAM111A, in addition to a ladder of higher molecular weight species (Figure 1B). Probing the same samples with SUMO1 and SUMO2 antibodies demonstrates that the higher molecular weight species are a combination of SUMO1 and SUMO2-modified FAM111A (Figure 1B).

Notably, overexpression of wild-type RNF4 strongly depletes the SUMO1 and SUMO2-modified species of FAM111A (Figure 1B). As expected, overexpression of RNF4^CS12^ enhances the higher molecular weight SUMO species of protease dead FAM111A (FAM111A^S541A^) however, it appears to deplete SUMOylated wild-type FAM111A (Figure 1B). This may be due to FAM111A and RNF4^CS12^ colocalization in subnuclear foci (see Figure 1E), wherein concentrated FAM111A protease activity could turnover SUMOylated FAM111A.

These data confirm that FAM111A is modified by SUMO1/2 and is a novel interactor of the human STUbL, RNF4. Taken together with high-throughput screens that identified proteins ubiquitinated and proteolytically regulated by RNF4 (Kumar et al., 2017), FAM111A is likely a novel target of STUbL-dependent regulation.

### Endogenous FAM111A Co-localizes with PCNA During Replication

FAM111A was identified associated with nascent chromatin (e.g. replication forks) in high-throughput studies (Alabert et al., 2014, Wessel et al., 2019). FAM111A also contains a PCNA interacting motif, which supports the colocalization of exogenously expressed FAM111A and PCNA (Alabert et al., 2014). The localization of endogenous FAM111A has been reported to be nucleolar, except during DNA replication when it was undetectable despite no change in protein abundance (Tarnita et al., 2019).

Using a validated commercial FAM111A antibody in multiple cell lines we detected the reported nucleolar localization, but also observed a focal nuclear pattern in some cells that co-localized with PCNA (Figure 1C). Based on the characteristic patterns of nuclear PCNA during DNA replication, and the high-throughput studies (Alabert et al., 2014, Wessel et al., 2019), our data confirm that FAM111A is associated with replication sites.

Interestingly, we also detected FAM111A at distinct subnuclear foci when U2OS cells were not in S phase, as judged by the lack of nuclear PCNA staining and the predominant nucleolar localization of FAM111A (Figure 1C). Such foci were not detected in the other cell lines tested. A key difference here is that U2OS cells maintain their telomeres via a recombination-based replication process called alternative lengthening of telomeres (ALT), as they do not have active telomerase (Bryan et al., 1997, Cesare and Reddel, 2010). ALT telomeres undergo DNA synthesis in the G2/M phases of the cell cycle at ALT-associated PML bodies (APBs) that contain PCNA amongst other DNA replication and repair factors (Dilley et al., 2016, Min et al., 2017, Pan et al., 2017, Zhang et al., 2019, Sobinoff and Pickett, 2020, Loe et al., 2020). Further analysis revealed that the atypical FAM111A foci in U2OS cells colocalize with PCNA and PML, as well as RNF4 (Figure 1C-E). Therefore, FAM111A localizes to sites of DNA synthesis during both bulk genome duplication, and the unscheduled DNA replication/repair events at ALT telomeres.

### Protease-Dependent Cytotoxicity of FAM111A

Genetic analysis of Kenney-Caffey syndrome 2 (KCS2) and osteocraniostenosis (OCS) revealed causative mutations in FAM111A (R569H and S342del, respectively; (Unger et al., 2013)). Based on genetic considerations these mutations were hypothesized to generate hyperactive variants of FAM111A (Unger et al., 2013). We therefore generated inducible cell lines to test the effects of overexpressing wild-type and mutant FAM111A.

Overexpression of wild-type FAM111A but not the protease dead mutant FAM111A^S541A^ inhibited cell growth (Figure 2A). Importantly, the KCS2 (FAM111A^R569H^) and OCS (FAM111A^S342del^) mutants were more potent inhibitors of cell growth than wild-type FAM111A, and ablating their protease activity with the *cis*-S541A mutation mitigated this toxicity (Figure 2A, **suppl. Fig. S1A**). Interestingly, HEK293T cells that constitutively express the SV40 large T antigen were relatively more resistant to wild-type, FAM111A^R569H^ or FAM111A^S342del^ overexpression than HEK293 cells (Figure 2A). This supports the suspected role of the SV40 Large T in inactivating FAM111A protease activity (see below, (Tarnita et al., 2019)).

**Figure 2.**
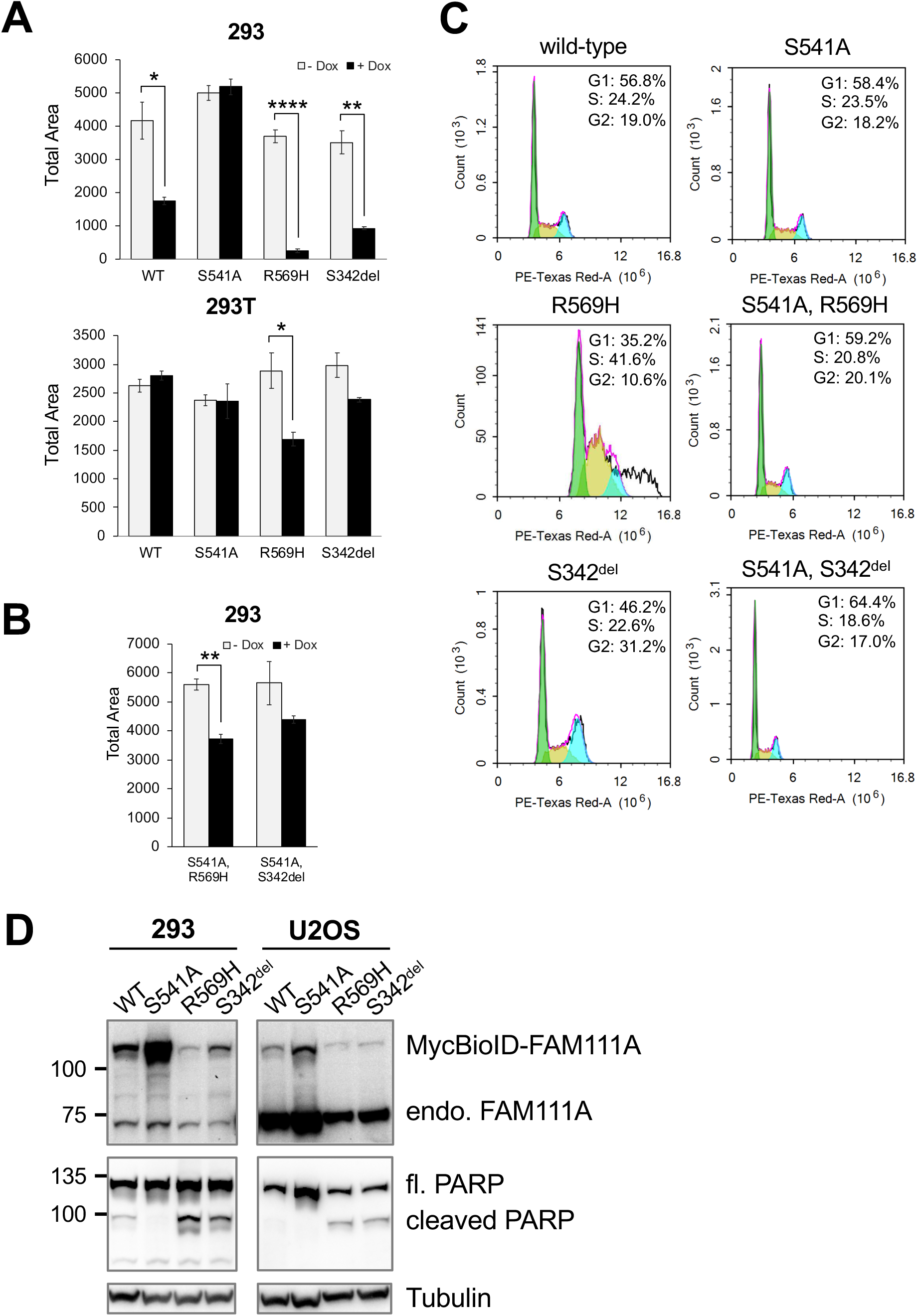
The effect of overexpressing wild-type or mutant FAM111A. **A, B.** HEK293, 293T/MycBioID-FAM111A cell lines were induced with doxycycline to express for 24 h. The colonies were fixed then stained with crystal violet. The cell growth was quantitated as total area on ImageJ. Values are mean ± s.d. of independent experiments (n = 3). \**p*<0.05, \*\**p*<0.01, \*\*\**p*<0.001, \*\*\*\**p*<0.0001 (two-tailed unpaired *t*-test). **C.** Cell-cycle profiling HEK293 FAM111A cell lines that were induced with doxycycline for 46 h and analyzed by flow cytometry. **D.** HEK293, U2OS/MycBioID-FAM111A cell lines were induced with doxycycline for 24 h. Total proteins from 105 cells were resolved by SDS-PAGE, and detected by Western blotting for both endogenous FAM111A (71 KDa) and MycBioID-FAM111A (110 KDa) as well as full length and cleaved PARP (116 and 89 KDa, respectively). The Tubulin Western blot was shown to compare sample loadings.

Overexpression of wild-type or FAM111A^S541A^ did not significantly perturb cell cycle profiles (Figure 2B, FACS -Dox in **suppl. Fig. S1B**). However, FAM111A^R569H^ caused a profound accumulation of cells in S phase, and FAM111A^S342del^ caused a marked increase in the G2/M population (Figure 2B). Given the localization of FAM111A to replication forks and its protease-dependent toxicity, these results suggest that the KCS2 and OCS patient mutants disrupt replication, possibly by unscheduled removal of replication factors.

Considering the above results, we tested if FAM111A overexpression can induce apoptosis. Cleavage of PARP is a well-characterized marker of caspase-dependent apoptosis (Soldani and Scovassi, 2002), and was weakly detected in cells expressing wild-type FAM111A but not protease-dead FAM111A^S541A^ (Figure 2C). Notably, PARP cleavage was stronger in cells expressing FAM111A^R569H^ or FAM111A^S342del^ than wild-type, and was abolished by the S541A mutation in *cis* (Figure 2C). Despite this, FAM111A^R569H^ and FAM111A^S342del^ expression was lower than wild-type FAM111A (Figure 2C). Also, exogenously expressed FAM111A was similar to the endogenous levels of FAM111A in HEK293 but much lower than those in U2OS, making their toxicity all the more striking (Figure 2C).

Interestingly, a FAM111A mutant lacking the SUMO2 modification sites experimentally mapped in a high-throughput study (FAM111A^K20,30,65R^, (Hendriks et al., 2017)) was expressed at similar levels to wild-type FAM111A but was more cytotoxic and induced higher levels of PARP cleavage (**suppl. Figs. S1A, C**). Mutation of an additional predicted (not experimentally mapped) SUMO acceptor lysine (FAM111A^K20,30,65,304R^) reduced FAM111A protein levels, cytotoxicity, and PARP cleavage proportionately (**suppl. Figs. S1A, C**).

Overall, our data are consistent with the prediction that KCS2 and OCS FAM111A patient mutants are gain-of-function (Unger et al., 2013), and that their protease activity is amplified with cytotoxic effects.

### Caspase-independent Effects of FAM111A Hyperactivity

We next tested the contribution of apoptotic induction to the profound cytotoxic effects of FAM111A^R569H^. To this end we used the well-characterized pan-caspase inhibitors z-VAD-fmk or Q-VD-OPh to reduce apoptotic signaling in cells overexpressing FAM111A, FAM111A^R569H^, or FAM111A^S541A^. As expected, pan-caspase inhibition blocked caspase-dependent PARP cleavage in both FAM111A and FAM111A^R569H^ expressing cells (Figure 3A). Despite this inhibition of apoptotic signaling, caspase inhibition provided no relief of FAM111A^R569H^ cytotoxicity (Figure 3B).

**Figure 3.**
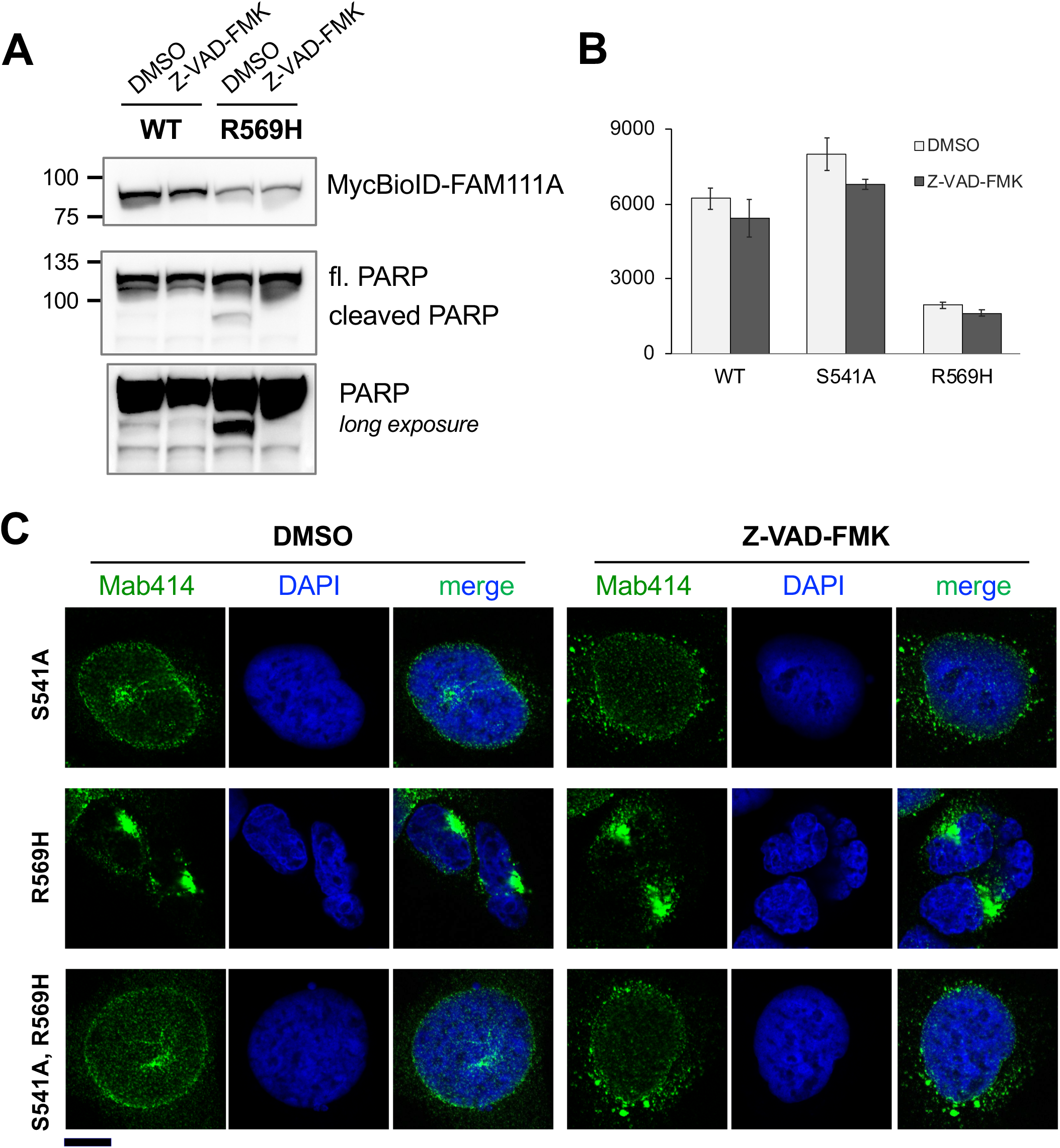
Caspase inhibitor Z-VAD-FMK inhibits PARP cleavage in cells overexpressing FAM111A^R569H^, but not cell death, nuclear shrinkage, and NPC redistribution. HEK293/MycBioID-FAM111A cell lines were treated with doxycycline for 22 h, along with DMSO or 100 µM Z-VAD-FMK were added to the cells (total inhibition of PARP cleavage was also observed when 50 µM Z-VAD-FMK was used). **A.** Total proteins from 105 cells were resolved by SDS-PAGE, and detected by Western blotting with antibodies against FAM111A or PARP. **B.** HEK293 cell lines were treated with doxycycline and Z-VAD-FMK for 24 h. The colonies were fixed then stained with crystal violet. The cell growth was quantitated as total area on ImageJ. Values are mean ± s.d. of independent experiments (n = 3). **C.** The cells were treated with doxycycline along with DMSO or Z-VAD-FMK for 24 h. They were fixed with formaldehyde, and stained with NPC antibody Mab414 and DAPI. The images were captured by confocal microscopy. Scale bar, 10 µm.

Growth inhibition by FAM111A^R569H^ could be related to its deregulation of S phase and cell cycle progression (Figure 2B) but this does not rule out additional effects of FAM111A on cell function. Therefore, we looked at the morphology and other features of cells expressing FAM111^R569H^. Overexpression of FAM111A^R569H^ caused the nuclei of most cells to “shrivel” and lose their characteristic smooth peripheries, resembling the appearance of early apoptotic nuclei (Figure 3C, **suppl. Fig. S2**). Also, nuclear pores underwent a dramatic redistribution, losing their typical granular appearance and “clumping” in regions of nuclear invagination or distortion (Figure 3C, **suppl. Fig. S2**). These phenotypes were not seen upon FAM111A^S541A^ overexpression and were dependent on the protease activity of FAM111A^R569H^ (Figure 3C, **suppl. Fig. S2**).

Notably, caspase inhibition again failed to block either of the above phenotypes caused by FAM111A^R569H^ expression (Figure 3C). Therefore, the protease activity of FAM111A^R569H^ appears to drive caspase-independent apoptosis-like phenotypes.

### Defining the FAM111A Proteomic Environment Identifies Candidate Targets

To begin to define candidate FAM111A protease targets driving the above morphological changes, we used proximity labeling (BioID) of FAM111A and FAM111A^S541A^ coupled to mass spectrometry-based protein identification (Roux et al., 2012). We reasoned that protease dead FAM111^S541A^ could generate enhanced or more durable biotin labeling of interactors than wild-type, similar to a substrate trap approach.

Candidate FAM111A interactors were identified from both asynchronous and S phase cells. Consistent with FAM111A’s co-localization with PCNA, several replication-associated proteins were identified including RFC1, RAD18, LIG1, and PCNA as enriched in S phase cells (Figure 4A).

**Figure 4.**
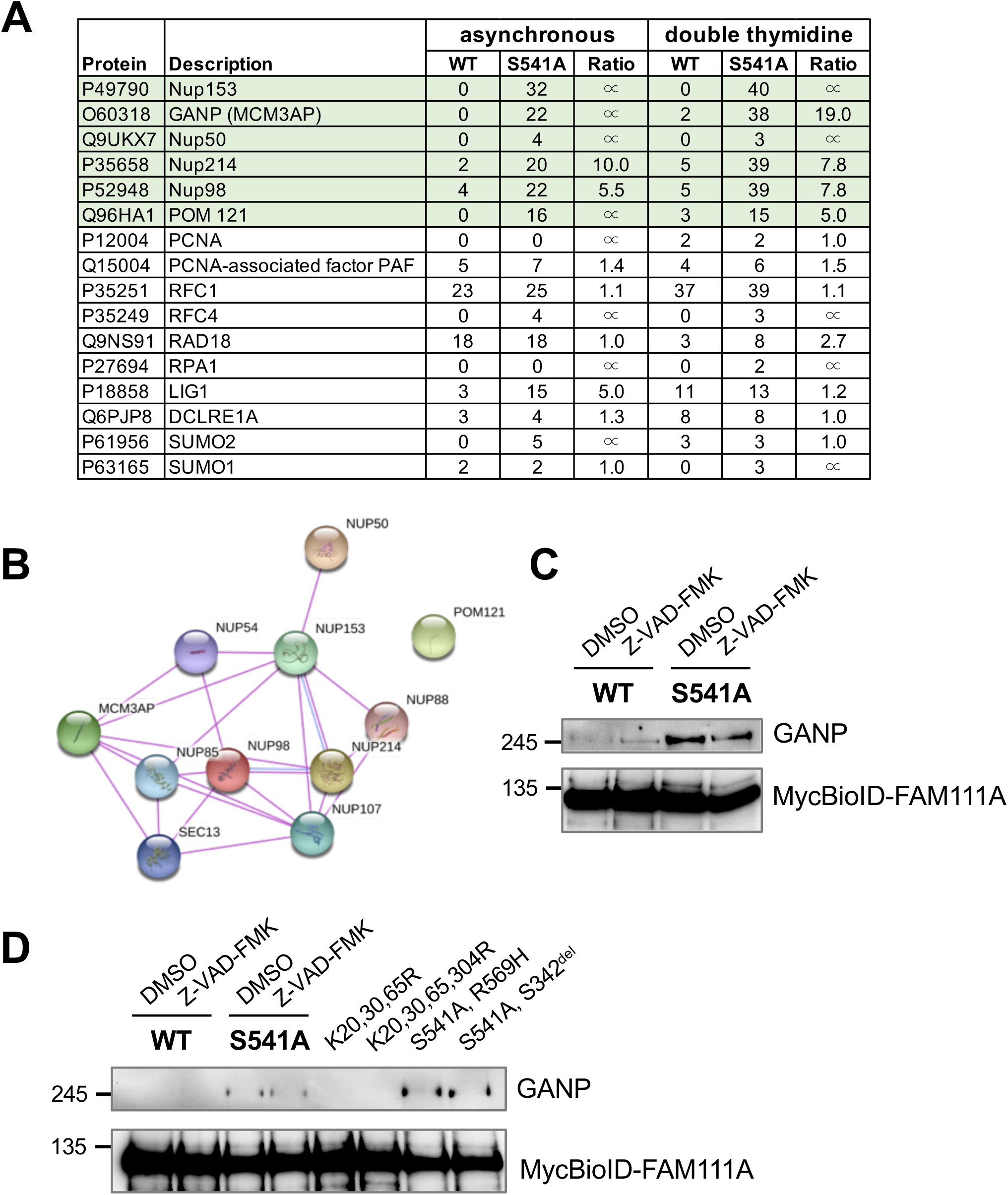
FAM111A BioID and mass spectrometry revealed interaction between FAM111A and NPC components. **A.** A subset of FAM111A interacting proteins identified by BioID screen. **B.** A STRING diagram showing the interaction of NPC components identified in FAM111A BioID screen. **C, D.** DMSO or Z-VAD-FMK treated HEK293/MycBioID-FAM111A cell lines were induced with doxycycline for 24 h. Total proteins from equal number of cells were pulled down using magnetic streptavidin beads, and analyzed by Western blotting using GANP and Myc antibodies.

One of the most highly enriched interactors identified in FAM111A^S541A^ over wild-type FAM111A purifications was the multi-functional mRNA export cofactor GANP, which is a scaffold of the TREX-2 complex at nuclear pores (Figure 4A; (Wickramasinghe et al., 2010)). In keeping with the localization of GANP, a number of nuclear pore proteins were also highly enriched such as NUP153, POM121, NUP214, NUP50 and NUP98 (Figure 4A & B).

To validate our FAM111A BioID results we used a GANP antibody to test its co-purification with FAM111A. Western blots revealed specific enrichment of GANP in FAM111A^S541A^ purifications but not those of wild-type FAM111A (Figure 4C). As FAM111A overexpression can induce apoptosis-like phenotypes, we tested if z-VAD-fmk affects the interaction between wild-type FAM111A and GANP, but found the results were unchanged (Figure 4C). We extended the analysis to include the FAM111A^R569H,S541A^ double mutant and the FAM111A^K20,30,65R^ SUMO site mutants. Again, GANP was only detected in BioID samples from FAM111A protease dead mutant expressing cells, including FAM111A^R569H,S541A^, indicating that this FAM111A patient mutation does not interfere with GANP interaction (Figure 4D).

Therefore, FAM111A^S541A^ has an increased duration proximal to GANP, and likely the other nuclear pore components. We have been unable to detect clear changes in the abundance of GANP or the nuclear pore complex components, but this does not exclude minor “clipping” activity by FAM111A that could influence the distribution or function of nuclear pores. Further analysis beyond the scope of this study is needed to define GANP and nucleoporins as proteolytic substrates of FAM111A, but there is a striking correlation with FAM111A’s function in controlling caspase-independent nuclear “permeability” during viral infection (see below).

### FAM111A Induces Changes in Nuclear Permeability during Viral Replication

FAM111A was originally identified as an SV40 host range restriction factor, which interacts with and is likely inactivated by the C-terminus of wild-type SV40 large T antigen (LT; (Fine et al., 2012)). Further studies indicated that FAM111A antagonizes the formation of replication centers of SV40 host range mutants containing LT C-terminal mutations such as SV40^HR684^ (Tarnita et al., 2019). Moreover, broadening the impact of FAM111A in viral restriction, it was also identified as a restriction factor for an orthopoxvirus host range mutant that lacks the serpin family serine protease inhibitor SPI-1 (Panda et al., 2017). Despite these advances, how FAM111A restricts viral challenges remains poorly defined.

Our results show that FAM111A activity can impair cellular viability and induce apoptosis-like phenotypes. However, the data so far are most relevant to the situation in KCS2 or OCS patients, where FAM111A is hyperactive (Unger et al., 2013). We therefore tested the impact of endogenous wild-type FAM111A on cells when challenged by SV40. To this end we transfected U2OS cells with wild-type SV40 or two host range mutants SV40^HR684^ and SV40^dl1066^ (Tarnita et al., 2019, Poulin and DeCaprio, 2006).

As previously reported, indirect immunofluorescence (IF) analysis of wild-type SV40 LT revealed progressive nuclear expression in U2OS cells over the 72 hr time course, and formation of defined replication centers in a small subset of these (Tarnita et al., 2019). In contrast, the SV40^HR684^ and SV40^dl1066^ host range mutants showed nuclear LT staining at 24 hr but over the next 48 hr the vast majority of transfected cells developed a pan-cellular LT distribution (Figure 5A, 5B, **suppl. Fig. S3**). In addition, nuclei appeared to partially shrivel and nucleoporins underwent a dramatic redistribution (**suppl. Fig. S4A**), as seen with FAM111A^R569H^ overexpression.

**Figure 5.**
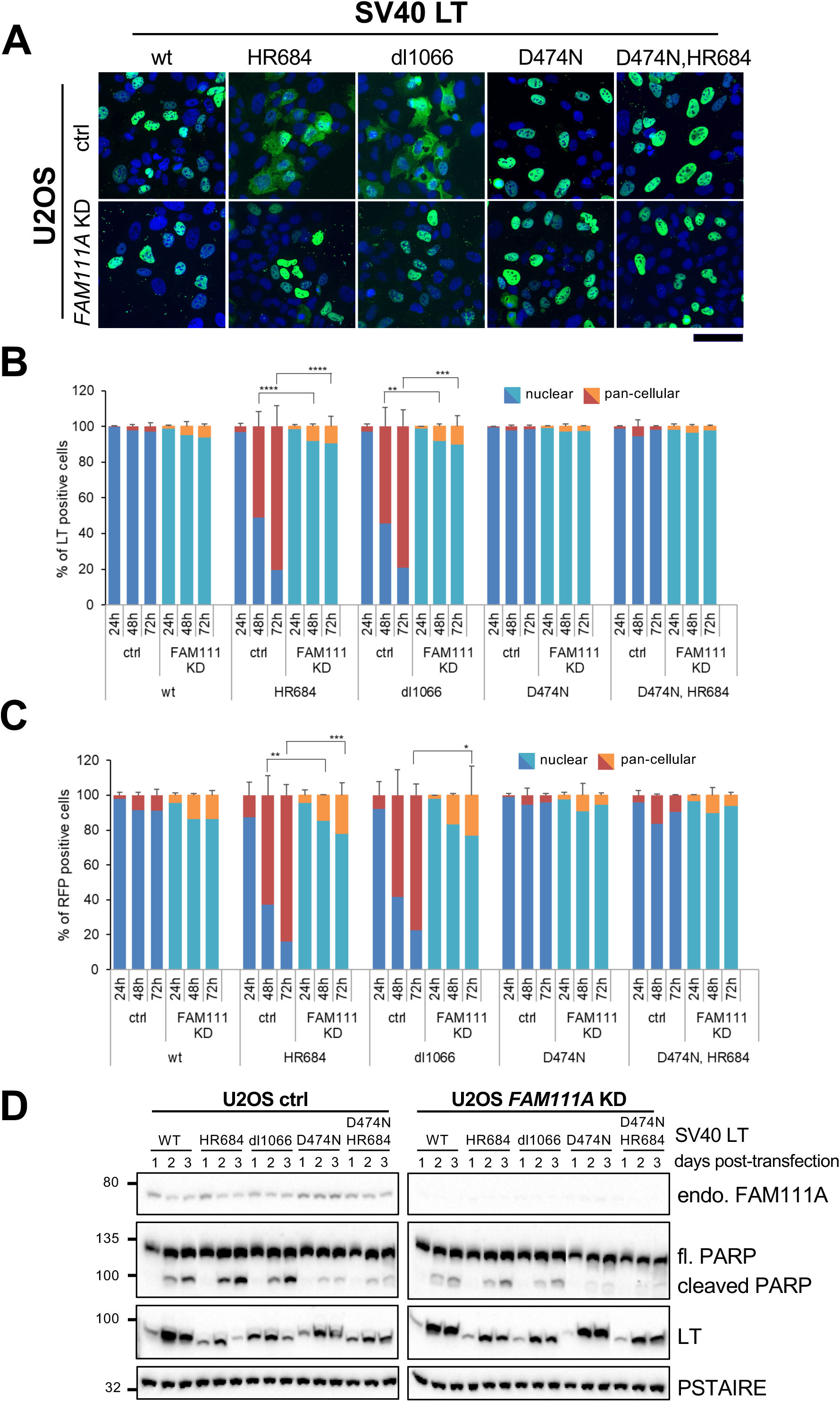
FAM111A affects nuclear barrier function during viral replication. **A.** Representative immunofluorescence images of SV40 LT localization 72 h after transfection into U2OS cells infected with control or FAM111A shRNA. Images were taken by Zeiss Axio Imager. LT is shown in green, and DAPI blue. Scale bar, 100 μm. **B.** Relative quantification of (**A**). Cells were fixed at indicated times post transfection. Mean values and standard deviations from the results of at least 3 independent experiments, minimum of 500 cells counted per condition. **C.** Relative quantification of RFP marker distribution upon co-transfection with SV40 LT variants into U2OS cells infected with control or FAM111A shRNA. Cells fixed at indicated times in at least two independent experiments, minimally 500 cells analyzed in each condition. In **B** & **C**, values are mean ± s.d. of independent experiments. \**p*<0.05, \*\**p*<0.01, \*\*\**p*<0.001, \*\*\*\**p*<0.0001 (two-tailed unpaired *t*-test). **D.** Western blot analysis of control or FAM111A-depleted U2OS cells transfected with SV40 LT variants. Cells were collected and lysed at indicated timepoints after SV40 transfection, immunoblotted with FAM111A, LT, PARP antibodies. Cdc2-PSTAIRE was used as a loading control.

To test if SV40^HR684^ and SV40^dl1066^ transfected cells actually lose normal nuclear barrier function, we co-expressed an independent 2XRFP-NLS nuclear reporter (Hatch et al., 2013). The loss of reporter compartmentalization mirrored that of the SV40 host range LT mutants (Figure 5C). Therefore, rather than a specific defect in the nuclear retention of LT host range mutants, nuclear barrier function is generally compromised. Importantly, depletion of FAM111A rescued the above phenotypes, including increased nuclear permeability (Figure 5A-C). Therefore, the wild-type but not SV40^HR684^ and SV40^dl1066^ LT antagonizes FAM111A activity, which otherwise causes nuclear dysfunction.

We next analyzed LT levels by western blot, as they reflect the ability of SV40 to transcribe and replicate its genome. As previously reported, wild-type SV40 LT expression peaked at 48 hr and sustained through 72 hr, whereas SV40^HR684^ and SV40^dl1066^ LT expression was weaker overall and tapered off by 72 hr (Figure 5D; (Tarnita et al., 2019)). Interestingly, PARP cleavage was higher in SV40^HR684^ and SV40^dl1066^ than wild-type SV40 transfected cells, being strongest at 72 hr when loss of nuclear barrier function is detected for the host range mutants (Figure 5D). Importantly, FAM111A depletion largely suppressed the differences in LT expression profiles of wild-type and host range mutant SV40, and reduced PARP cleavage in SV40^HR684^ and SV40^dl1066^ cells (Figure 5D).

FAM111A appears to execute its restrictive function during viral DNA replication (Tarnita et al., 2019, Panda et al., 2017), which fits with its localization to cellular DNA replication sites and the negative effects of FAM111A^R569H^ on processive replication. To determine if SV40 replication triggers the FAM111A-dependent loss of nuclear barrier function, we used an LT mutation (D474N) that abolishes its helicase activity and SV40 genome replication (Huang et al., 2010).

We generated the single SV40^D474N^ LT mutant and the double SV40^D474N,HR684^ LT mutant for analysis by IF and western blot. SV40^D474N^ LT was almost exclusively nuclear at all time points, and western analysis revealed a wild-type expression profile that did not taper at 72 hr (Figure 5A-D). Importantly, the SV40^D474N,HR684^ double mutant behaved like the SV40^D474N^ single mutant in terms of nuclear localization and expression profile (Figure 5A-D).

In contrast to the SV40^HR684^ LT, FAM111A depletion had no impact on either the nuclear localization or expression profile of SV40^D474N^ or SV40^D474N,HR684^ LT. Thus, it is not simply SV40^HR684^ LT expression that causes nuclear permeability changes, rather; it is SV40 replication in the presence of FAM111A activity.

## Discussion

We find that FAM111A exhibits protease-dependent cytotoxicity, which is amplified by the KCS2 and OCS patient mutants. This likely explains the dominant effects of FAM111A^R569H^ and FAM111A^S342del^ over wild-type in the heterozygous disease state (Unger et al., 2013). Our results are consistent with recent data showing that FAM111A patient mutants have higher intrinsic protease activity, undergoing autocleavage *in vitro* more strongly than wild-type (Kojima et al., 2020).

Importantly, we show that hyperactive FAM111A negatively impacts nuclear integrity, S phase progression, and cell viability, effects that could underlie KCS2 and OCS. For example, parathyroid development (amongst other systems) could be impaired by hyperactive FAM111A, leading to a number of KCS2 and OCS phenotypes such as hypoparathyroidism, hypocalcemia, and associated skeletal dysplasia (Unger et al., 2013).

Intriguingly, phenotypes caused by FAM111A hyperactivity such as nuclear deformation, “clumping” of nuclear pores, and substrate detachment, are insensitive to pan-caspase inhibitors (despite robust inhibition of PARP cleavage). At a minimum, even if residual caspase activity were present, we would expect mitigation of some of the above phenotypes, which is not the case. In this regard, there have been reports of a caspase-independent loss of nuclear function(s) in early apoptosis similar to what we report, which is instead dependent on an unidentified serine protease (Egger et al., 2003, O’Connell and Stenson-Cox, 2007, Ferrando-May et al., 2001, Strasser et al., 2012, Kopeina et al., 2018). In the future, it will be interesting to determine if FAM111A is the serine protease in question.

We also analyzed the role of FAM111A in restricting the replication of SV40 host range (HR) mutants (Tarnita et al., 2019, Fine et al., 2012). Our results are broadly consistent with the published studies in that FAM111A prevents SV40 HR mutant replication in restrictive cells. Interestingly however, we found that FAM111A causes the loss of normal nuclear structure, pore distribution and barrier function when SV40 HR mutants attempt to replicate. These phenotypes are strikingly similar to those caused by the KCS2 and OCS FAM111A patient mutants, and could explain how FAM111A restricts SV40 HR mutant replication.

Unlike restrictive cells such as U2OS, HEK293 cells are permissive for SV40 replication, do not lose nuclear barrier function over time, and have abundant replication centers for both wild-type and HR mutant SV40. Interestingly, we found that endogenous FAM111A is much less abundant in HEK293 cells versus U2OS and other cells types tested (**suppl. Fig. S4B**). Therefore, based on our data, HEK293 cells could in part be permissive hosts for SV40 due to low endogenous FAM111A levels.

Although SV40 HR mutants induce slightly more PARP cleavage than wild-type SV40 in restrictive cells, the difference is inconsistent with the dramatic loss of nuclear barrier function caused by SV40 HR mutants (~80%). Instead, the increase in cells with pan-cellular LT over time (>72 hr) in the absence of late apoptotic phenotypes indicates that FAM111A triggers nuclear barrier dysfunction to derail SV40 HR mutant replication, which can cause delayed apoptosis in a subset of cells.

As mentioned above, an unidentified serine protease was implicated in the caspase-independent loss of nuclear barrier function in response to different types of cell stress (Egger et al., 2003, O’Connell and Stenson-Cox, 2007, Ferrando-May et al., 2001, Strasser et al., 2012, Kopeina et al., 2018). In addition, early studies revealed that RPV and VACV host range mutants that lack the serine protease inhibitor SPI-1 cause alterations in the nuclear morphology of restrictive cells (Brooks et al., 1995, Shisler et al., 1999). Together with the fact that FAM111A is now a known host restriction factor for VACV SPI-1∆ (Panda et al., 2017), our data suggest that the disruption of cell function by nuclear barrier loss could be a common function for FAM111A in restricting viral replication.

We identified a number of nuclear pore components as candidate targets of FAM111A, and confirmed that GANP more stably associates with protease dead over wild-type FAM111A. Interestingly, whereas FAM111A is enriched on nascent chromatin at the replication fork, several nucleoporins and GANP are actually *depleted* (Wessel et al., 2019, Alabert et al., 2014). Nucleoporins and associated factors such as GANP make chromatin contacts to spatially organize chromatin, control its outputs such as transcription, and maintain genome stability (Ibarra and Hetzer, 2015).

It is tempting to speculate that FAM111A protease activity could help displace nuclear pore-chromatin interactions to facilitate replication fork progression. Indeed, actively transcribed genes are transiently released from the nuclear periphery during replication (Brickner and Brickner, 2011). Also, a recent study found that cells lacking FAM111A have elevated levels of Top1 cleavage complexes (Top1cc) (Kojima et al., 2020). Rather than a direct role in Top1cc repair, FAM111A could reduce replication-transcription collisions by displacing GANP etc., which otherwise indirectly lead to increased Top1cc. In future, it will be important to determine if FAM111A hyperactivity due to KCS2/OCS mutations or viral challenge causes excessive nucleoporin and GANP processing, which leads to the observed loss of nuclear function.

Consistent with its localization to replication forks, we also identified a number of replication factors in close proximity to FAM111A such as RFC1, PCNA, RAD18, and LIG1. However, unlike GANP and the nucleoporins these factors were identified equally with wild-type or protease dead FAM111A baits, suggesting they may not be targets. Instead, the PCNA loader RFC1 collaborates with FAM111A to restrict poxvirus replication (Panda et al., 2017). Also, restriction of poxvirus (RPXV∆SPI-1) and polyomavirus (SV40^HR684^) host range mutants occurs upon attempted viral genome replication (Figure 5 & (Tarnita et al., 2019, Brooks et al., 1995, Luttge and Moyer, 2005)).

We favor a model in which the loading of PCNA acts as a DNA sensor that recruits FAM111A to endogenous replication sites or viruses that utilize/attract PCNA (Sowd and Fanning, 2012, Postigo et al., 2017). During unscheduled viral DNA replication or in KCS2/OCS patients, FAM111A is hyperactivated and cytotoxic due to the absence of normal protease (auto) regulation. Wild-type SV40 LT and VACV SPI-1 are able to inhibit FAM111A protease activity, preventing nuclear dysfunction and enabling viral replication. In the future, it will be interesting to determine endogenous regulatory mechanisms for FAM111A protease activity, and how it is tolerated at ALT telomeres that are replicated outside S phase.

## Materials and Methods

### Cell culture

Human embryonic kidney cell lines 293 and 293T, human osteosarcoma cell line U2OS, human colon cancer line HCT116, and human cervical adenocarcinoma cell line HeLa were obtained from American Type Culture Collection (ATCC). Human epithelial cells immortalized with hTERT cell line RPE was obtained from Denchi lab (Loe et al, 2020). These cell lines were cultured in Dulbecco’s modified Eagle’s medium (DMEM) supplemented with 10% fetal bovine serum (FBS).

Unless otherwise indicated, the following drug concentrations were used: doxycycline (1 µg/ml in H_2_O, Sigma-Aldrich), biotin (50 µM in H_2_O, Sigma-Aldrich, B4501), Z-VAD-FMK (50 or 100 µM in DMSO, APExBIO, A1902), Q-VD-OPh (20 µM in DMSO, Sigma-Aldrich, SML0063).

### Plasmids

Full-length human *RNF4*, *FAM111A* cDNAs were PCR amplified and inserted into destination vectors using In-Fusion seamless cloning (Takara Bio). The pTRIPz-MycBioID vector allows for doxycycline-inducible expression of RNF4 or FAM111A with N-terminal fusion of a Myc tag and a promiscuous biotin ligase (MycBioID) (Roux et al., 2012). Point mutations in RNF4, FAM111A, and SV40 large T antigen (LT) were introduced using the QuikChange Site-Directed Mutagenesis kit (Agilent Technologies) according to the manufacturer’s protocol. All cloning and mutagenesis were verified by sequencing. Plasmids used in this study are listed in supplemental table (Table S1).

### RNAi

Stable RNAi was achieved by viral shRNA generated in pLKO.1-blast (Bryant et al., 2010). Knockdown was verified by standard western blot, normalizing to Tubulin or Cdc2 expression. pLKO.1 lentiviruses were constructed according to the Addgene pLKO.1 protocol (www.addgene.org), and target sequences were based Mission shRNA database (Sigma-Aldrich). RNAi target sequences used in this research include:

Control-sh: 5’-CCTAAGGTTAAGTCGCCCTCG-3’

FAM111A-sh1: 5’-GTCAATGTGTAAGGGTGACAT-3’

### Transfection, virus production and transduction

For lentivirus production, pTRIPz or pLKO.1 plasmid was co-transfected with lentiviral Packaging vectors (pMD2.G and psPAX2) into HEK293T using TransIT-LT1 (Mirus) transfection reagent following the manufacturer instruction. All supernatants were passed through 0.22 μm to remove cell debris. The supernatants were either immediately applied to the target cells, or frozen in liquid nitrogen for further use. For lentivirus transduction (pLKO.1, pTRIPz), subconfluent U2OS, HEK293, or 293T cultures, the day after plating, were infected with virus-containing supernatants for 12–16 h at 37°C. Viral supernatants were then replaced with fresh growth medium, cultured for a further 72 h with appropriate antibiotic selection. Puromycin (0.5 µg/ml for U2OS; 2 µg/ml for HEK293, 293T), and blasticidin (12 µg/ml) were used. To generate stable cell lines (Table S2), the transduced cells were cultured in the presence of antibiotic selection until all control cells were dead. SV40 plasmid transfections were conducted using Trans-IT LT1 (Mirus) transfection reagent following the manufacturer’s instructions.

### Immunofluorescence and microscopy

The cells grown on coverslips were fixed in 4% formaldehyde in PBS for 10 min, then permeabilized with PBS containing 0.2% Triton X-100 for 5 min, at room temperature. The cells stained with PCNA were pre-extracted in Triton X-100 buffer containing 0.5% Triton X-100, 20 mM Hepes-KOH (pH 7.9), 50 mM NaCl, 3 mM MgCl2, and 300 mM Sucrose for 2 min on ice, then fixed in 3% formaldehyde, 2% sucrose in PBS for 10 min at room temperature. The cells were incubated with primary antibodies diluted in blocking solution: 5% Normal Goat Serum (Biolegend, 927502), or 0.2% (w/v) cold water fish gelatin (Sigma, G-7765) and 0.5% (w/v) BSA (Sigma, A-2153) in PBS for 1 h at room temperature or at 4 oC overnight. Following staining with secondary antibodies and 0.1 µg/ml 4’,6-diamidino-2-phenylindole dihydrochloride (DAPI, Sigma, D-9542) diluted in 5% NGS-PBS, cells were mounted onto glass slides using ProLong Gold Antifade (Invitrogen #P36934). Primary antibodies used for immunofluorescence included: FAM111A (Abcam, ab184572 (1:100)), SV40LT (Abcam, ab16879 (1:250)), PCNA (Santa Cruz, sc-56 (1:200)), PML (Santa Cruz sc-966 (1:200)), anti-Nuclear Pore Complex proteins (Mab414, Abcam, ab24609 (1:200)). Secondary antibodies used were Alexa Fluor Secondary Antibodies (Life Technologies).

Confocal mages were acquired with a Zeiss LSM 880 inverted spectral imaging confocal microscope (Carl Zeiss Microscopy) equipped with 63x/1.4 oil immersion high NA objective lens, using standard settings in Zen microscope software. Data was exported and processed using ImageJ software (NIH). Digital images were also taken using Zeiss Axio Imager M1 with 20x and 60x oil immersion objectives.

### Clonogenic survival assays

HEK293, 293T cells were counted. Five hundred cells were seeded in six well plates, allowed to adhere overnight, and treated next day with 1 ug/ml of doxycycline. Twenty-four hours later, doxycycline was removed, and fresh medium was added. Cells were cultured for another four days until visible colonies have formed, then fixed and stained with 0.5% crystal violet. The plates were scanned on LI-COR (LI-COR Biosciences), and cell growth was quantitated as total area using ImageJ (NIH).

### Flow cytometry

For analysis of DNA content, cells were fixed with 70% EtOH, and stained with 10 µg/ml propidium iodide, 100 µg/ml RNase A in PBS containing 0.1% Triton X-100. Flow cytometry analysis was performed on a NovoCyte 3000 Flow Cytometer (ACEA Biosciences) using NovoExpress software.

### BioID labeling and affinity purification of biotinylated proteins

Cells were incubated for 24 h in complete media supplemented with doxycycline and biotin. After washing with PBS, cells (2 to 5 × 10^6^ cells for small-scale analysis; 2 × 10^7^ cells for large scale analysis) were collected and frozen. Cell pellets were lysed and processed as described (Roux et al., 2012). Briefly, each 10^6^ cells were suspended in 100 µl of strong lysis buffer containing 50 mM Tris, pH 7.5, 500 mM NaCl, 0.4% SDS, 5 mM EDTA, 1 mM DTT, 1x Halt Protease Inhibitor Cocktail (ThermoFisher, 78430), and 1 mM PMSF. The lysate was sonicated in a Misonix waterbath Sonicator. Triton X-100 was added to 2% final concentration, and sonication was repeated. Equal volume of cold 50 mM Tris, pH 7.4 was added, followed by additional sonication. After centrifugation at 16,000 × *g* for 10 min at 4 oC, supernatants were incubated with Dynabeads MyOne Streptavidin C1 (ThermoFisher, 65002) at 4 oC for 2 h (20 µl of beads were used for small-scale analysis; 100 µl for large scale preparation for mass spectrometry).

Afterward, the beads were washed at 25°C sequentially in 1 ml of 2% SDS (wash buffer 1) twice, once each in 0.1% deoxycholate, 1% Triton X-100, 500 mM NaCl, 1 mM EDTA, and 50 mM Hepes, pH 7.5 (wash buffer 2), and in 250 mM LiCl, 0.5% NP-40, 0.5% deoxycholate, 1 mM EDTA, and 10 mM Tris, pH 8.1 (wash buffer 3), and finally twice in 50 mM Tris, pH 7.4, 50 mM NaCl (wash buffer 4). For Western blot analysis, proteins were eluted from the magnetic beads with 35 µl of 2x NuPAGE LDS Sample Loading Buffer (ThermoFisher, NP0007) with 100 mM DTT at 100 °C for 5 min. For mass spectrometry analysis, the beads were washed and stored in 100 ml of 8M urea, 100 mM Tris (pH 8.5) until further processing.

### Western blotting

Cells were lysed in strong lysis buffer described in the section above. The whole cell lysate was combined with NuPAGE LDS Sample Loading Buffer and 100 mM DTT, separated by SDS-PAGE and transferred to nitrocellulose. Immunoblotting was performed as described (Nie et al., 2017). Antibodies used for immunoblotting included: FAM111A (Abcam, ab184572 (1:1000)), Myc (9E10, Scripps Antibody Core Facility (1:3000)), PARP (Cell Signaling, 9542 (1:1000)), GANP (Bethyl, A303-128A (1:1000)), SV40LT (Abcam, ab16879 (1:3000)), tubulin alpha (Sigma, T9026 (1:10,000)), and Cdc2-PSTAIR (Abcam, ab10345 (1:15,000)). Following incubation with HRP-conjugated secondary antibodies, proteins were detected using SuperSignal West Dura Extended Duration Substrate (ThermoFisher, 34076) on the ChemiDox XRS+ System (BioRad).

### Protein identification by mass spectrometry

Protein samples were reduced and alkylated by sequential incubation with 5mM Tris (2-carboxyethyl) phosphine for 20 minutes at room temperature and 10 mM iodoacetamide reagent in the dark at room temperature for 20 additional minutes. Proteins were digested sequentially at 37 °C with lys-C for 4 hours followed by trypsin for 12 hours. After quenching the digest by the addition of formic acid to 5% (v/v), peptides were desalted using Pierce C18 tips (Thermo Fisher Scientific), dried by vacuum centrifugation, and resuspended in 5% formic acid. Peptides were fractionated online using reversed phase chromatography on in-house packed C18 columns. The 140 minute gradient of increasing acetonitrile was delivered using a Dionex Ultimate 3000 UHPLC system (Thermo Fisher Scientific). Peptides were electrosprayed into the mass spectrometer by the application of a distal *2.2 kV* spray voltage. MS/MS data were acquired using an Orbitrap Fusion Lumos mass spectrometer operating in Data-Dependent Acquisition (DDA) mode consisting of a full MS1 scan to identify peptide precursors that were subsequently targeted by MS2 scans (Resolution = 15,000) using high energy collision dissociation for the remainder of the 3 second cycle time. Data analysis was performed using the Integrated Proteomics bioinformatic pipeline 2 (Integrated Proteomics Applications, San Diego, CA). Database searching was performed using the ProLuCID algorithm against the EMBL Human reference proteome (UP000005640 9606). Peptides identifications were filtered using a 1% FDR as estimated using a decoy database. Proteins were considered present in a sample if they had 2 or more unique peptides mapping to them. Relative comparisons between samples to identify candidate FAM111A interacting proteins was done using raw peptide spectral counts.

## Supporting information

Proteomic Table

## Acknowledgements

We thank Dr. James DeCaprio for the SV40 genome plasmids. We are grateful to Taylor Loe and the Scripps Research community for their support and helpful discussions. This study was supported by NIH grants GM068608 and GM122987 awarded to MNB.

## Author Contributions

MNB, MN, MO, ELD, YJA, and JAW devised and executed experiments in support of the study. MN and MO performed most of the presented experiments. YJA and JAW carried out mass spectrometry-based protein identification analysis of our RNF4 and FAM111A BioID samples. All authors contributed to manuscript writing and figure preparation.

