## Supplementary Figures and Legends for "Activation of FAM111A Protease Induces Defects in Nuclear Function that Likely Underlie its Roles in Disease and Viral Restriction"

**This file includes:**

Supplementary Figures S1-S4
Supplementary Tables S1 and S2

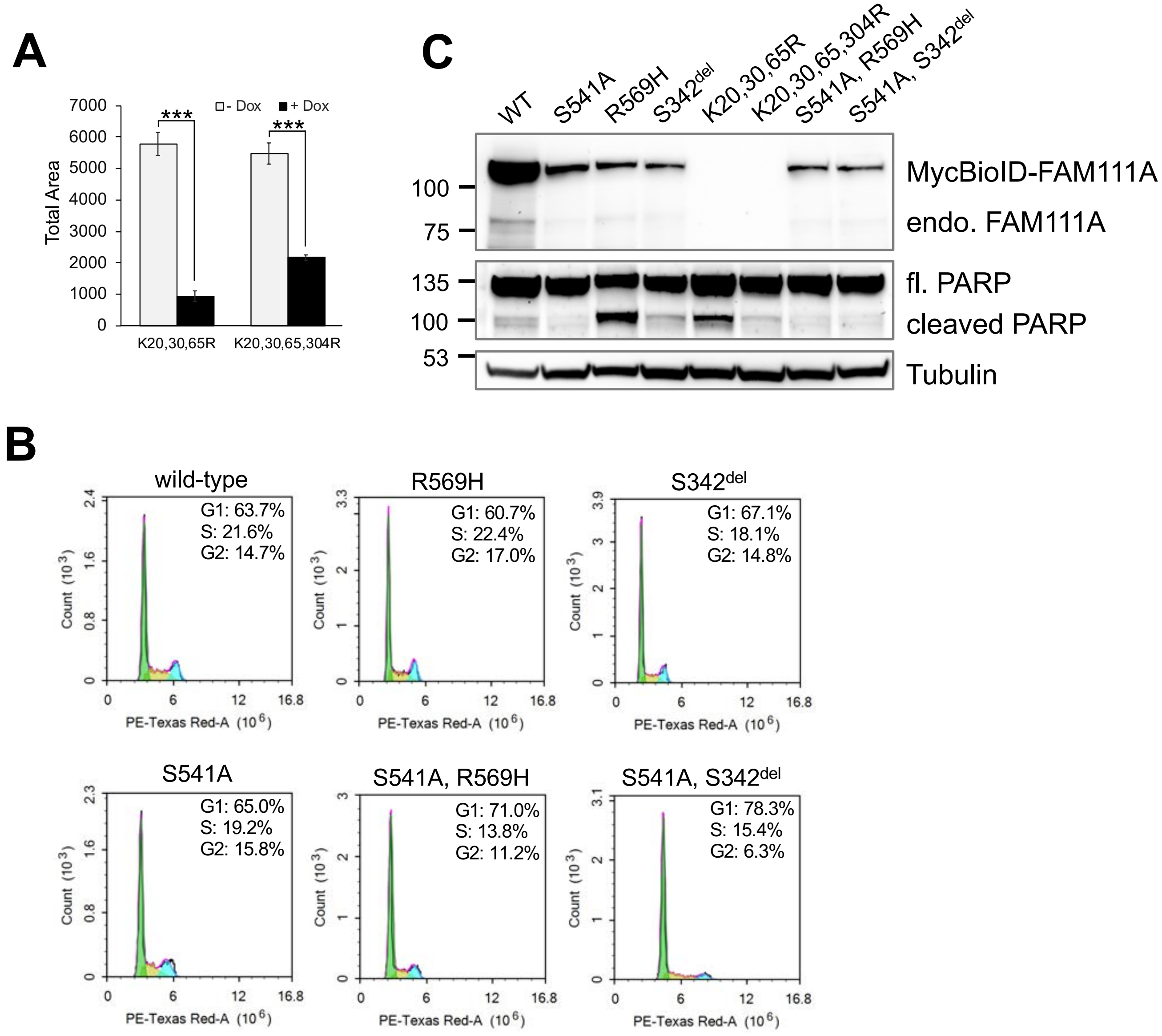


**Supplementary Figure S1**. **A.** HEK293/MycBioID-FAM111A cell lines were induced with doxycycline to express for 48 h. The colonies were fixed then stained with crystal violet. The cell growth was quantitated as total area on ImageJ. Values are mean ± s.d. of independent experiments (n = 3). ***p*<0.01, ****p*<0.001 (two-tailed unpaired *t*-test). **B.** Flow cytometry analysis of cell-cycle profiles of HEK293 FAM111A cell lines seeded and fixed as in Figure 2A, but not treated with doxycycline. **C.** HEK293/MycBioID-FAM111A cell lines were induced with doxycycline for 24 h. Total proteins from $10^5$ cells were analyzed by Western blotting for both endogenous FAM111A (71 KDa), MycBioID-FAM111A (110 KDa), full length and cleaved PARP (116 and 89 KDa, respectively), as well as Tubulin levels.

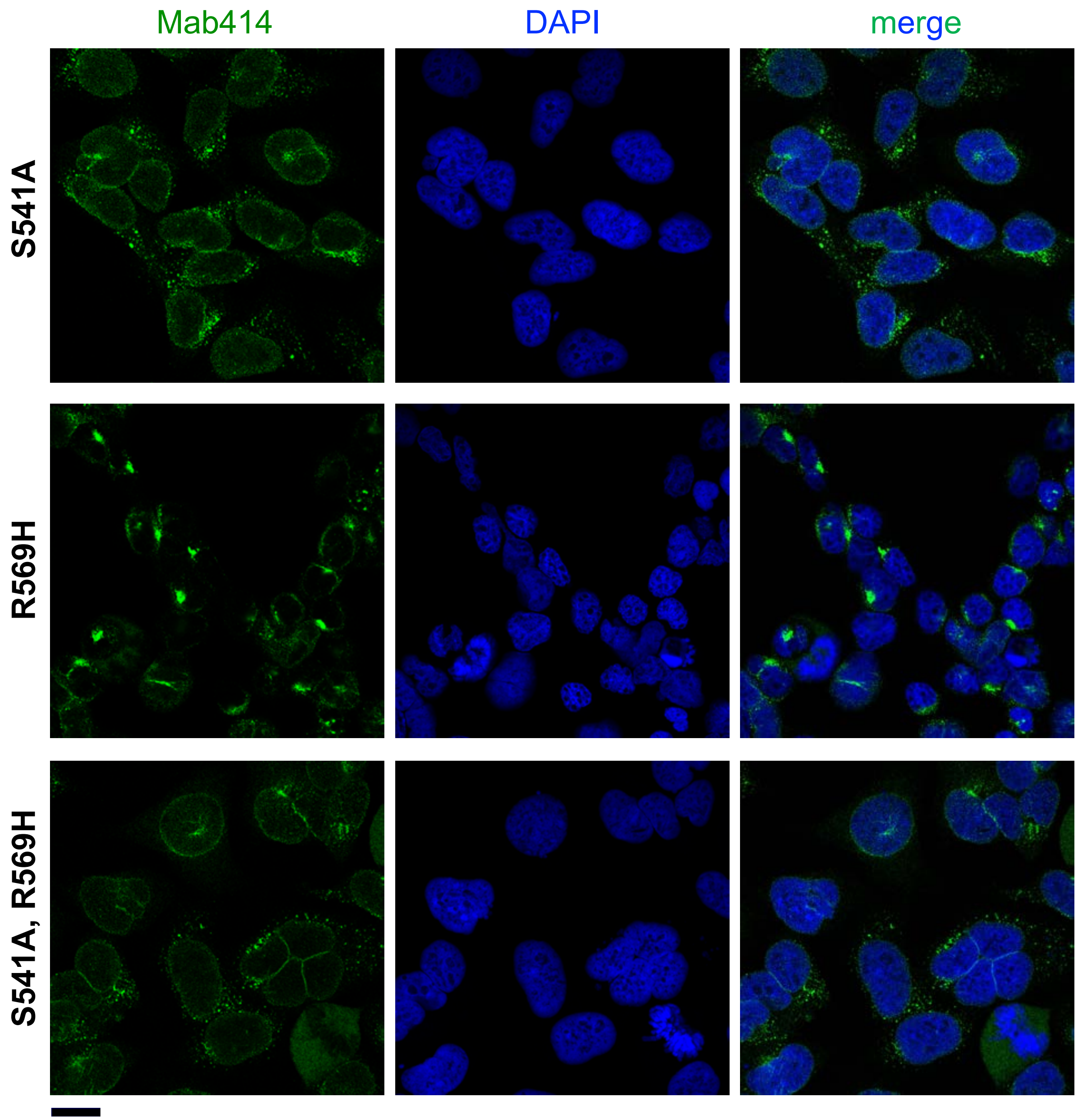


**Supplementary Figure S2.** The fields of cells captured through a 63x/1.4 objective of LSM880 confocal microscope (selection of individual cells is shown in Figure 3C). Scale bar, 20 μm.

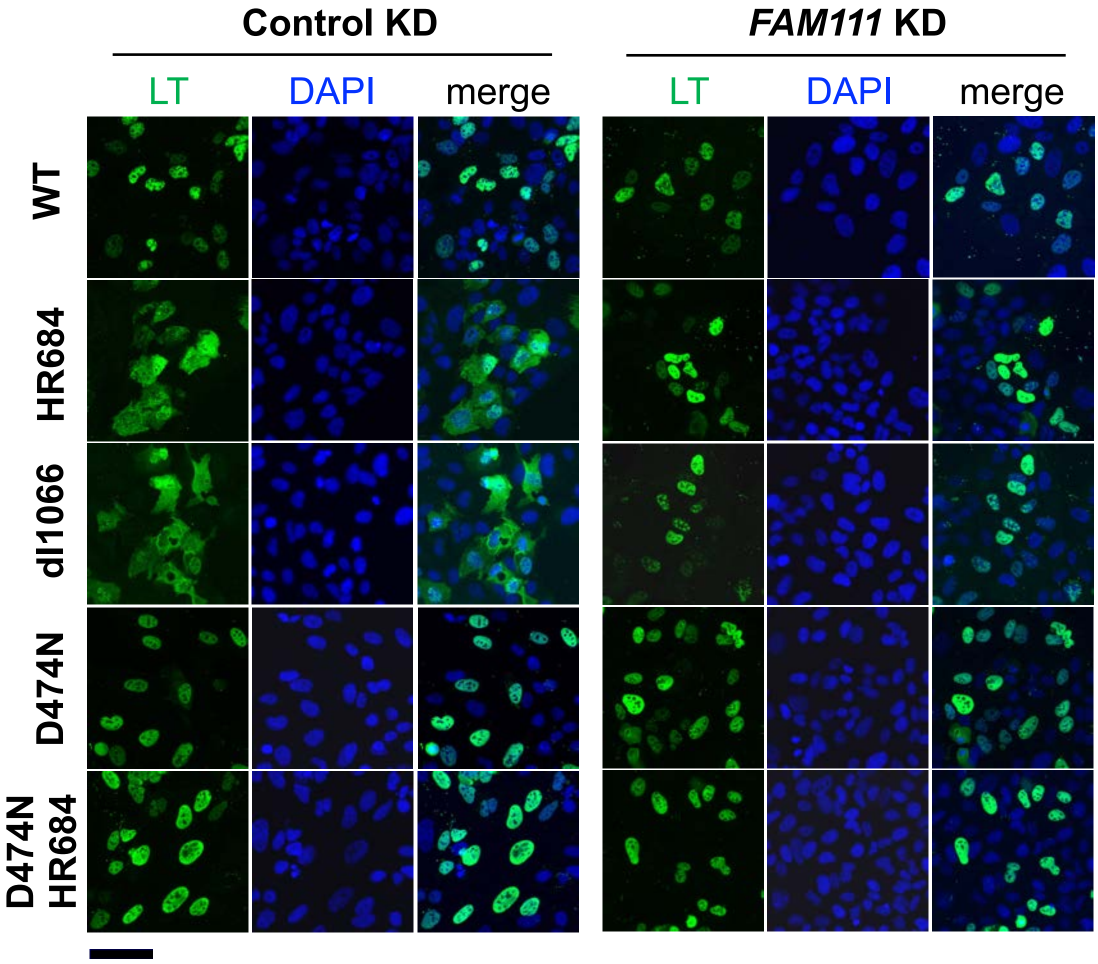


**Supplementary Figure S3.** Representative immunofluorescence images of SV40 LT (green) and DAPI (blue) staining in control or FAM111A shRNA infected U2OS cells 72hr after transfection with SV40 plasmids. Images were taken by Axio Imager 20x/0.6 objective. Scale bar, 100 μm.

**A**

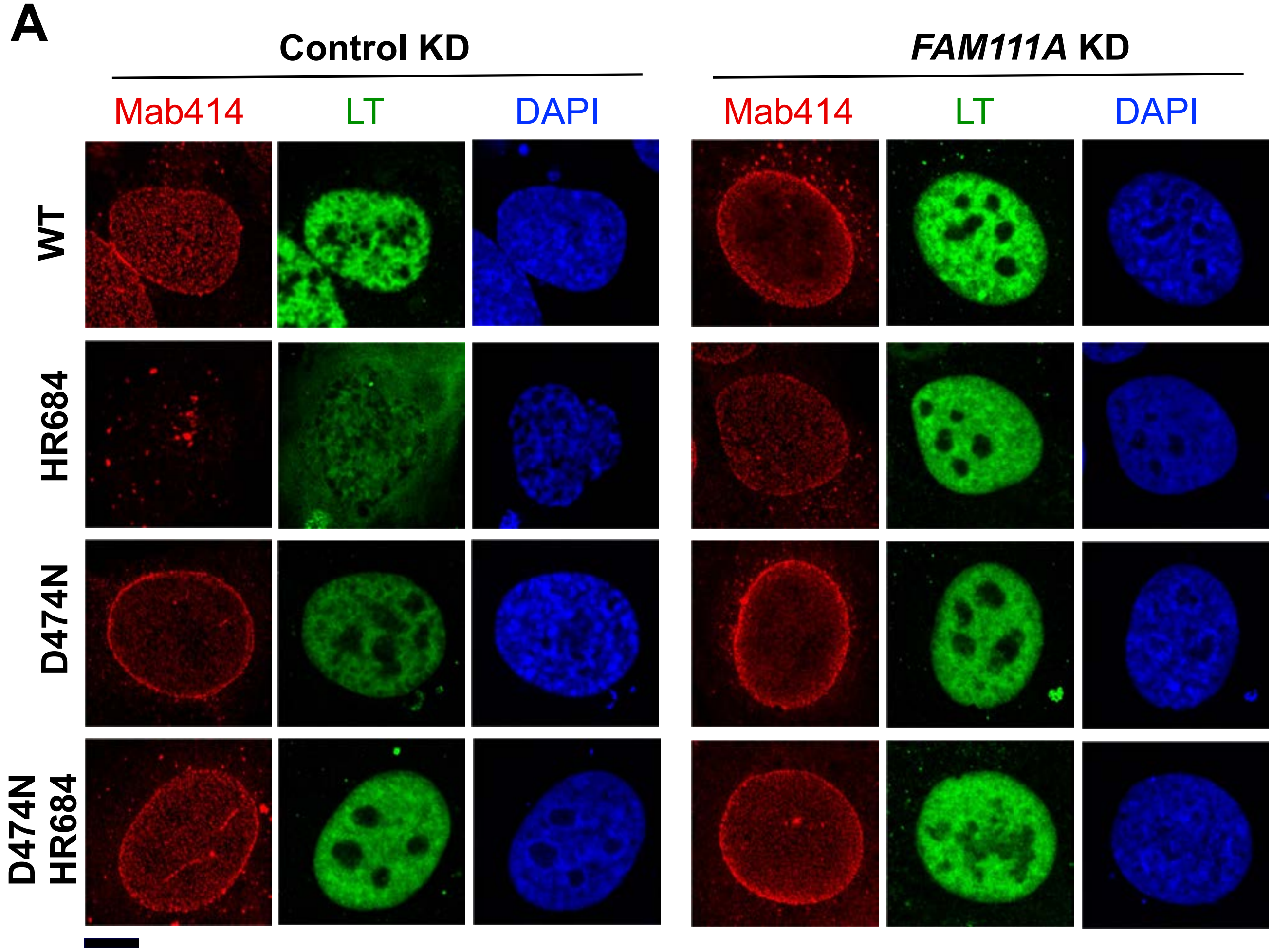


**B**

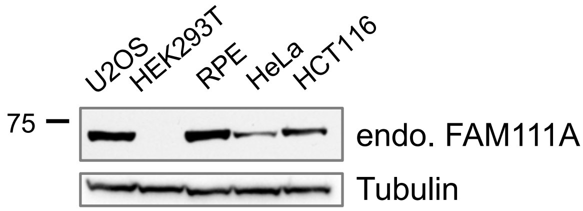


**Supplementary Figure S4. A.** Representative immunofluorescence images of SV40 LT and NPC (Mab414) localization 72 h after transfection into U2OS cells infected with control or FAM111A shRNA. Images were captured through a 63x/1.4 objective of LSM880 confocal microscope. Scale bar, 10 μm. **B.** Western blot analysis of FAM111A levels in cell lines. Total proteins from equal number of cells were loaded in each lane. The blots were probed with FAM111A and Tubulin antibodies.

**Supplementary Table S1.** List of plasmids

| Plasmid # | Plasmid Name | Reference |
|---|---|---|
| pNB171 | pMD2.G | Addgene (#12259) |
| pNB172 | psPAX2 | Addgene (#12260) |
| pNB173 | pLKO-1-Blast-scrambled-sh | this study |
| pNB210 | pTRIPz-(M)YFP-Eed | Addgene (#82512) |
| pNB252 | pLPC-MycBioID-RNF4 | this study |
| pNB265 | pLPC-MycBioID-RNF4-CS12 | this study |
| pNB268 | pTRIPz-MycBioID-RNF4-CS12 | this study |
| pNB317 | pEGFP-C3-FAM111A | this study |
| pNB371 | pBS-SV40-WT | Tarnita et al. 2019 |
| pNB372 | pBS-SV40-HR684 | Tarnita et al. 2019 |
| pNB373 | pBS-SV40-dl1066 | Tarnita et al. 2019 |
| pNB349 | pTRIPz-MycBioID-FAM111A | this study |
| pNB350 | pTRIPz-MycBioID-FAM111A-S541A | this study |
| pNB392 | pTRIPz-MycBioID-FAM111A-R569H | this study |
| pNB393 | pTRIPz-MycBioID-FAM111A-S342del | this study |
| pNB398 | pTRIPz-MycBioID-FAM111A-K20,30,65R | this study |
| pNB400 | pTRIPz-MycBioID-FAM111A-K20,30,65,304R | this study |
| pNB401 | pTRIPz-MycBioID-FAM111A-S541A-R569H | this study |
| pNB403 | pTRIPz-MycBioID-FAM111A-S342del-S541A | this study |
| pNB408 | pDEST-mCherry-NLS-tag-RFP | Hatch et al., 2013 |
| pNB417 | pBS-SV40-D474N | this study |
| pNB418 | pBS-SV40-D474N-HR684 | this study |

**Supplementary Table S1.** Stable cell lines

| Cell Type | Expression Plasmid |
|---|---|
| U2OS | pTRIPz-MycBioID-FAM111A |
| U2OS | pTRIPz-MycBioID-FAM111A-S541A |
| U2OS | pTRIPz-MycBioID-FAM111A-R569H |
| U2OS | pTRIPz-MycBioID-FAM111A-S342del |
| U2OS | pTRIPz-MycBioID-RNF4-CS12 |
| HEK293 | pTRIPz-MycBioID-FAM111A |
| HEK293 | pTRIPz-MycBioID-FAM111A-S541A |
| HEK293 | pTRIPz-MycBioID-FAM111A-R569H |
| HEK293 | pTRIPz-MycBioID-FAM111A-S342del |
| HEK293 | pTRIPz-MycBioID-FAM111A-K20,30,65R |
| HEK293 | pTRIPz-MycBioID-FAM111A-K20,30,65,304R |
| HEK293 | pTRIPz-MycBioID-FAM111A-S541A-R569H |
| HEK293 | pTRIPz-MycBioID-FAM111A-S342del-S541A |
| HEK293T | pTRIPz-MycBioID-FAM111A |
| HEK293T | pTRIPz-MycBioID-FAM111A-S541A |
| HEK293T | pTRIPz-MycBioID-FAM111A-R569H |
| HEK293T | pTRIPz-MycBioID-FAM111A-S342del |
